# Optimal feedback mechanisms for regulating cell numbers

**DOI:** 10.1101/292920

**Authors:** Saurabh Modi, Abhyudai Singh

**Affiliations:** Department of Biomedical Engineering, University of Delaware, Newark, DE, USA.; Department of Electrical and Computer Engineering, University of Delaware, Newark, DE, USA.; Department of Mathematical Sciences, University of Delaware, Newark, DE, USA.

## Abstract

How living cells employ counting mechanisms to regulate their numbers or density is a long-standing problem in developmental biology that ties directly with organism or tissue size. Diverse cells types have been shown to regulate their numbers via secretion of factors in the extracellular space. These factors act as a proxy for the number of cells and function to reduce cellular proliferation rates creating a negative feedback. It is desirable that the production rate of such factors be kept as low as possible to minimize energy costs and detection by predators. Here we formulate a stochastic model of cell proliferation with feedback control via a secreted extracellular factor. Our results show that while low levels of feedback minimizes random fluctuations in cell numbers around a given set point, high levels of feedback amplify Poisson fluctuations in secreted-factor copy numbers. This trade-off results in an optimal feedback strength, and sets a fundamental limit to noise suppression in cell numbers. Intriguingly, this fundamental limit depends additively on two variables: relative half-life of the secreted factor with respect to the cell proliferation rate, and the average number of factors secreted in a cell’s lifespan. We further expand the model to consider external disturbances in key physiological parameters, such as, proliferation and factor synthesis rates. Intriguingly, while negative feedback effectively mitigates disturbances in the proliferation rate, it amplifies disturbances in the synthesis rate. In summary, these results provide unique insights into the functioning of feedback-based counting mechanisms, and apply to organisms ranging from unicellular prokaryotes and eukaryotes to human cells.

## 1 Introduction

In order to achieve physiological function various animals regulate their morphology [1]. For example, tissue size may be regulated by either controlling the sizes of individual cells, or the number of constituent cells [2–6]. How individual cells maintain size homeostasis has been extensively studied across organisms ranging from bacteria, animal and plant cells [7–16]. Interestingly, data reveals cell-autonomous control strategies that regulate cellular growth or timing of cell-cycle events to suppress aberrant deviations in cell size around an optimal size specific to that cell type [17–20]. In contrast, mechanisms that control cell numbers have been less understood.

One strategy to control cell numbers is for cells to secrete a molecule that accumulates in the extracellular space, and is sensed by other cells in the population [21–23]. Binding of this secreted factor to cell surface receptors activates signaling pathway that inhibit cell proliferation, thus creating a negative feedback loop (Fig. 1). Such feedback control via secreted factors has been reported in many organisms including *Myxococcus xanthus* [24], *Vibrio fischeri* [25], *Bacillus subtilis* [26], *Dictyostelium discoideum* [27, 28], multicellular animals [29, 30] and plants [31]. For illustration purposes, we show some published data on *Dictyostelium discoideum* in Fig. 1, where the proliferation rates decrease monotonically with increasing buildup of the secreted factor.

**Figure 1:**
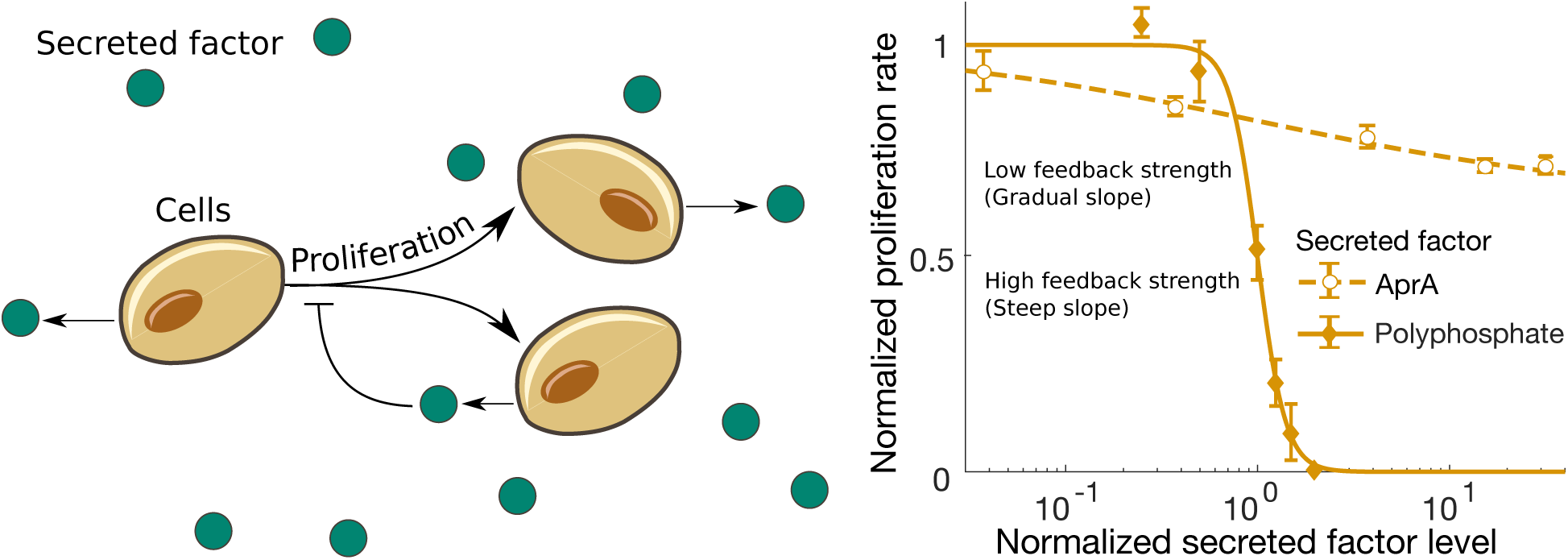
Controlling cell numbers by feedback control through secreted extracellular factors. *Left.* Schematic of a cell population, where each cell proliferates and secretes a molecule in the extracellular space. The buildup of secreted factors is sensed by other cells in the population. The factor functions to inhibit cell proliferation creating homeostasis in cell numbers. *Right.* Data showing proliferation rates of *Dictyostelium discoideum* cells with respect to the levels of two different secreted factors (*AprA* and *Polyphosphate*). While the inhibition of proliferation rate by *AprA* is gradual, that by *Polyphosphate* is relatively steep and occurs over a narrower range of factor levels. The error bars are the standard errors corresponding to the raw data, and lines represent Hill equations fitted to the data. We refer the reader to [32, 33] for further biological details and experimental procedures.

The feedback strategy illustrated in Fig. 1 creates an interesting tradeoff, where cells incur a cost to produce the factor and would prefer to keep its synthesis to as low as possible. However, a side effect of low levels is shot noise or Poisson fluctuations in secreted factor copy numbers that are propagated to cell numbers. Here we study this tradeoff through a mathematical model that incorporates three different noise mechanisms: stochastic proliferation of cells; stochastic synthesis of the secreted factor from single cells; and external disturbances in model parameters. Our analysis reveals that depending on the source of noise, negative feedback can either buffer or amplify random fluctuations in cell numbers around a given set point. Moreover, when multiple noise sources are present, then an optimal feedback strength provides the most efficient noise buffering. This optimal feedback sets up a fundamental lower limit for minimizing fluctuations in cell numbers, and we systematically study how this limit scales with different parameters. We start by formally introducing the mathematical model followed by its stochastic analysis.

## 2 Stochastic model formulation

Let ***x***(*t*) and ***z***(*t*) denote the number of cells, and the number of secreted factors at time *t*, respectively. Cells are assumed to proliferate exponentially with a rate *g* (***z***), with *g* being a monotonically decreasing function as illustrated in Fig. 1. Moreover, cells are removed (or die) from the population at a rate *γ*_***x***_. Finally, each cell synthesizes and secretes the factor at a rate *k*_***z***_, and these secreted factors decay with a rate *γ*_***z***_. The stochastic formation of this model is shown in Table 1 and consists of four probabilistic events that increase/decrease the population counts by one. The propensity functions in the last column determine how often the events occur. For example, the propensity function for the cell proliferation event is *g* (***z***) ***x*** which implies that the probability this event will occur in the next time interval (*t, t* + *dt*) is *g* (***z***) *xdt*, and whenever the event occurs the cell count increases by one.

**Table 1:** Stochastic model of cell proliferation and feedback inhibition via secreted factors.

| Probabilistic event | Change in population counts | Propensity function |
| --- | --- | --- |
| Cell proliferation | $\mathbf{x}(t) \mapsto \mathbf{x}(t) + 1$ | $g(\mathbf{z}(t)) \mathbf{x}(t)$ |
| Cell death | $\mathbf{x}(t) \mapsto \mathbf{x}(t) - 1$ | $\gamma_{\mathbf{x}} \mathbf{x}(t)$ |
| Secreted factor synthesis | $\mathbf{z}(t) \mapsto \mathbf{z}(t) + 1$ | $k_{\mathbf{z}} \mathbf{x}(t)$ |
| Secreted factor degradation | $\mathbf{z}(t) \mapsto \mathbf{z}(t) - 1$ | $\gamma_{\mathbf{z}} \mathbf{z}(t)$ |

Throughout the paper we denote by 〈x(*t*)〉 as the mean value of the stochastic process ***x***(*t*), and by 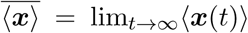 its steady-state value. Based on the stochastic model in Table 1, the population averages evolve as

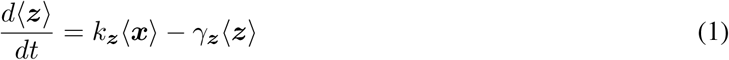

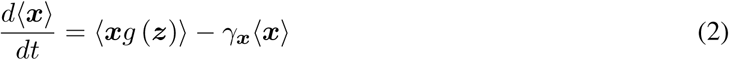

[34, 35]. In the deterministic mean-field limit where

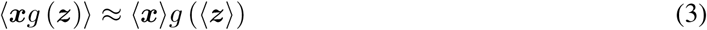

we obtain the following approximated nonlinear system

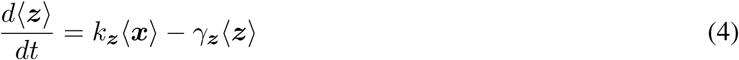

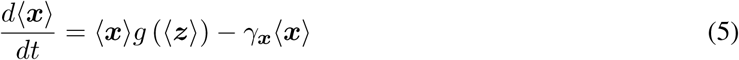

that has a unique equilibrium defined by the solution to the equations

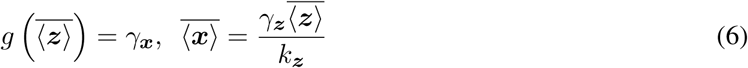

as long as *g*(0) > *γ*_***x***_ > *g* (*∞*). Having determined the mean equilibrium population counts, we next quantify the negative feedback strength by the dimensionless parameter

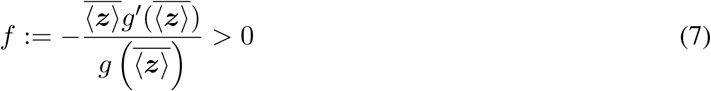

that is essentially the log sensitivity of *g* at 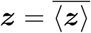. Local stability analysis of the equilibrium point yields the following eigenvalues of the linearized system

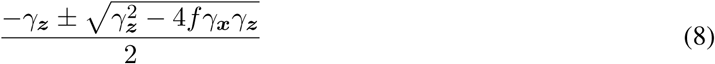

and shows that the equilibrium of the nonlinear system (4)-(5) is stable for *f* > 0. Note that sufficiently strong negative feedback (*f* > *γ*_***z***_/4*γ*_***x***_) results in complex eigenvalues with negative real parts, and in this case as the noise kicks the system out of equilibrium, the relaxation dynamics will be oscillatory.

## 3 Optimal feedback strength for regulating cell numbers

A stochastic simulation of the feedback system via the Gillespie algorithm reveals an intriguing feature - while low levels of feedback strength attenuate random fluctuations in cell numbers, a strong negative feedback amplifies fluctuations (Fig. 2). We further investigate this effect by developing approximate analytical formulas for the noise in cell numbers, where noise is quantified by the steady-state coefficient of variation of ***x***(*t*). A well-known problem that often arises when dealing with nonlinear stochastic systems is unclosed moment dynamics - time evolution of lower order moments depends on higher order moments [37]. While a number of closure methods have been developed to tackle this issue [38–51], we circumvent this problem by exploiting the Linear Noise Approximation (LNA) [52]. Assuming small fluctuations in ***x***(*t*) and ***z***(*t*) around their respective steady-state means, LNA works by linearizing the nonlinear propensity functions

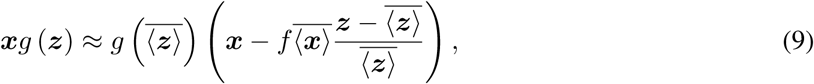

and with this approximation all propensity functions in Table 1 are now linear functions of the state space. Time evolution of statistical moments is obtained using the following result from our prior work [35, 37]: the time derivative of the expected value of an arbitrary function *ψ*(***x**, **z***) is given by

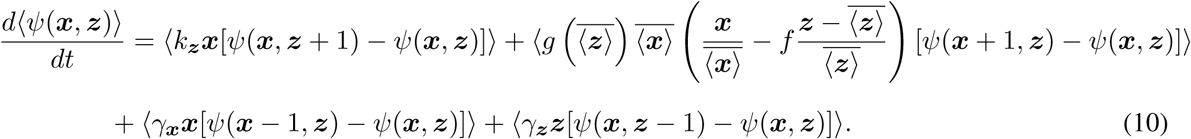

Substituting appropriate monomials for *ψ*(***x**, **z***) above yields the following system of differential equations for all the first and second order moments of ***x***(*t*) and ***z***(*t*)

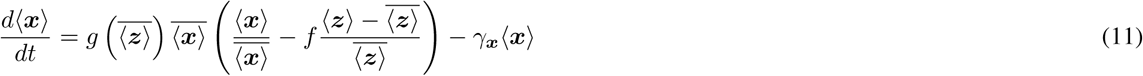

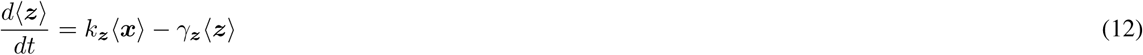

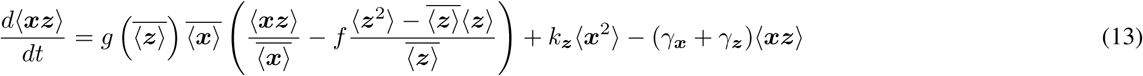

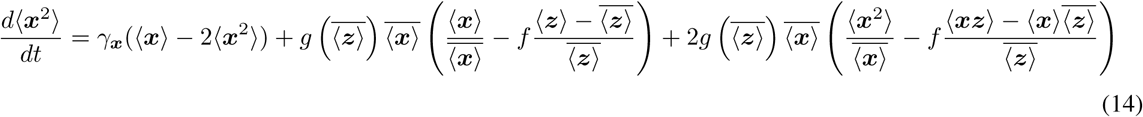

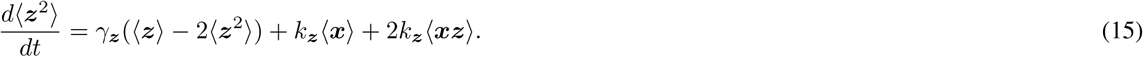

Solving the moment equations at steady state we obtain the following noise (coefficient of variation squared) in cell numbers

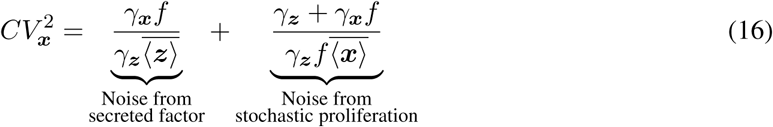

which can be decomposed into two terms. The first term represents the noise contribution from Poisson fluctuations in ***z***(*t*) (noise from secreted factor) and the other term is the contribution from stochastic proliferation/death of cells. Note that *CV**_x_*** decreases as the secreted factor half-life decreases, which makes intuitive sense as then the regulator tracks the cell numbers more faithfully. Interestingly, the first term is amplified, and second term is attenuated with increasing feedback strength *f* (Fig. 2). These opposing effects result in 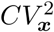 being minimized at an optimal feedback strength1

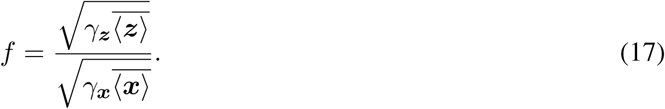

Replacing (17) in (16) and using (3) yields the fundamental limit of noise suppression in cell numbers as

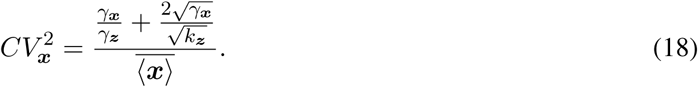

**Figure 2:**
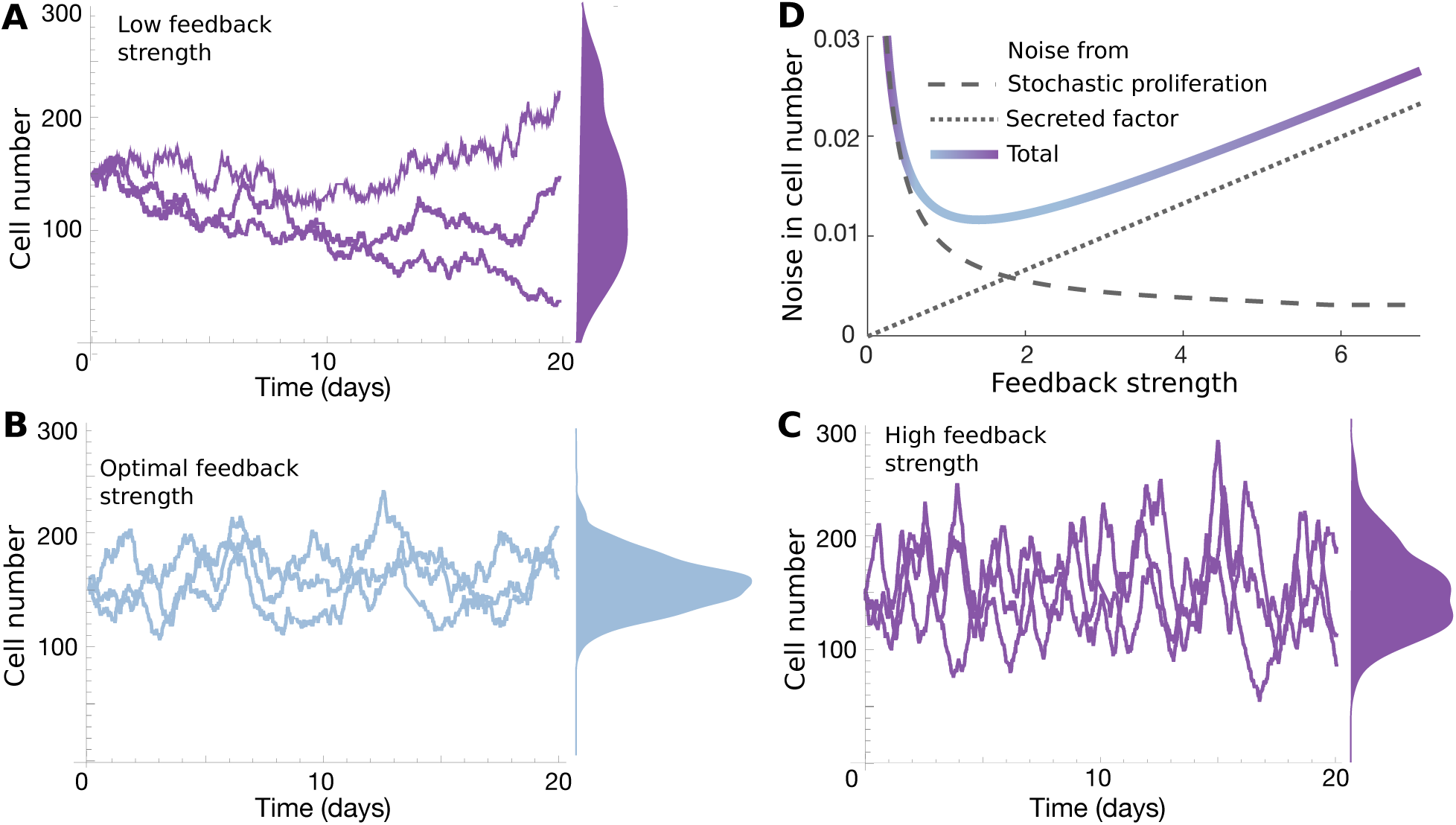
Noise in cell numbers is minimized at an intermediate level of negative feedback. Stochastic simulation of the model presented in Table 1 using [36] for different feedback strengths A.*f* = 0.01 (Low), B. *f* = 0.6 (Optimal), C. *f* = 10 (High). For a given mean number of cells, the spread in cell numbers first reduces, and then increases with increasing feedback strength. D. Plotting the total noise, and the different noise components in (16) as a function of *f*. While the noise contribution from stochastic proliferation reduces, the contribution from stochastic secreted-factor synthesis and decay increases creating a U-shape profile of the total noise. Parameters chosen for all the above subfigures were: *γ*_*z*_ = 3*γ*_*x*_ = 3 day^−1^, 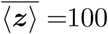, 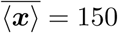.

While inverse scaling of 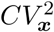 with respect to the mean 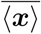 is reminiscent of Poisson-like fluctuations, the numerator value can be made small by having a short-lived secreted factor (high *γ*_*****z*****_) *and* a high factor synthesis rate *k*_***z***_. It is interesting to point out that the ratio *N* = *k*_***z***_/*γ*_***x***_ is the average number of secreted factors made by an individual cell in its lifespan, and the fundamental noise limit scales as 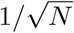, but the scaling is 1/*γ*_***z***_ with respect to the factor decay rate.

## 4 Incorporating external disturbances in physiological parameters

Our analysis up till now assume constant model parameters, and we now expand the model to allow for external disturbances that transform parameters into stochastic processes.

### 4.1 External disturbance in the proliferation rate

Data shows that the cellular growth rate can randomly fluctuate with some memory over many generations [53–57], and its not hard to imagine these features carrying over to the cell proliferation rate. Motivated by these findings we assume that the proliferation rate *g* (***z**, **y***) is affected by an external factor ***y*** whose stochastic dynamics is modeled as an Ornstein-Uhlenbeck process

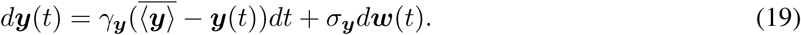

Here, 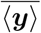 is the mean level of external disturbance, ***w***(*t*) is the Wiener process, *σ*_***y***_ and *γ*_***y***_ are parameters that represent the strength of noise and the time-scale of fluctuations in ***y***, respectively. Linearizing the proliferation rate

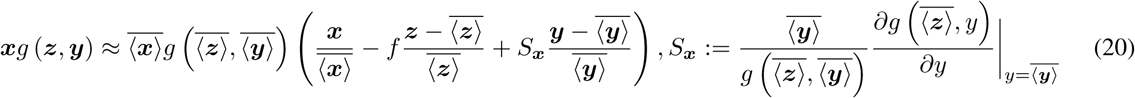

where *S*_***x***_ is the log sensitivity of the proliferation rate to the external disturbance. With this approximations we again repeat the analysis of obtaining moment equations and solving it to get noise in cell numbers. We write the moment dynamics of an arbitrary function *ψ*(***x**, **y**, **z***) using the following result

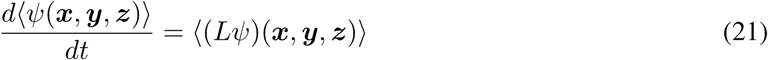

[34], where

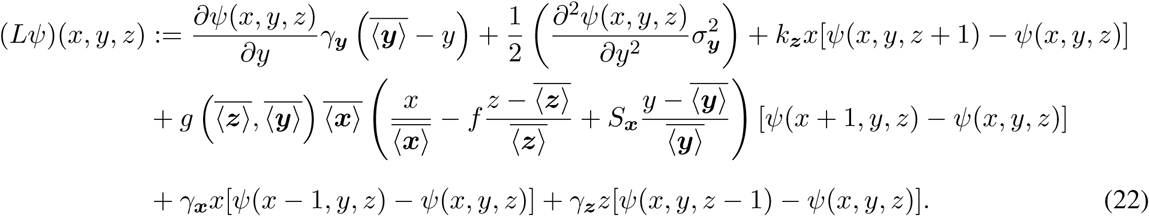

Substituting appropriately monomials for *ψ*(***x**, **y**, **z***) we obtain the following moment dynamics

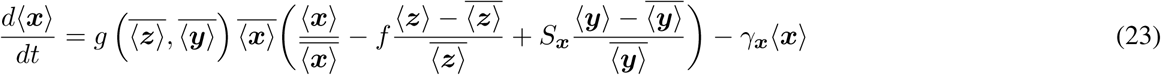

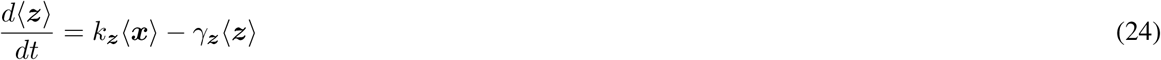

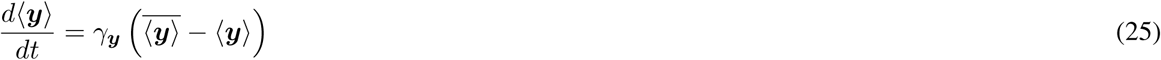

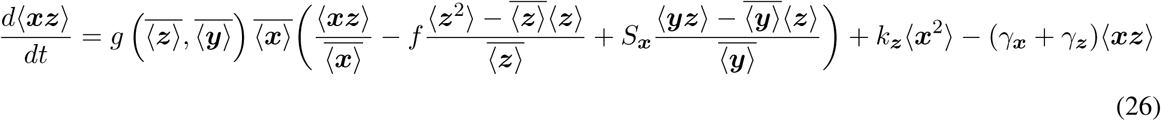

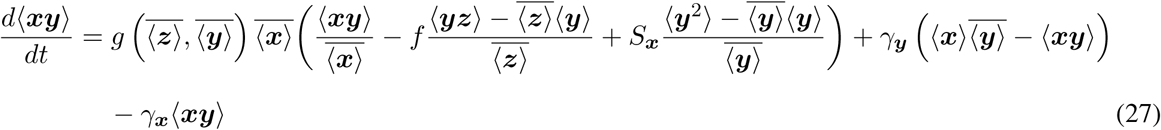

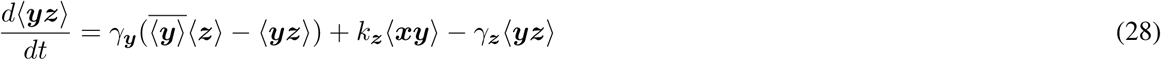

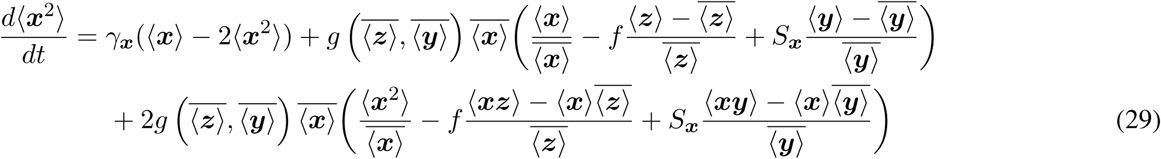

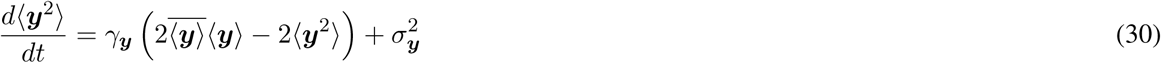

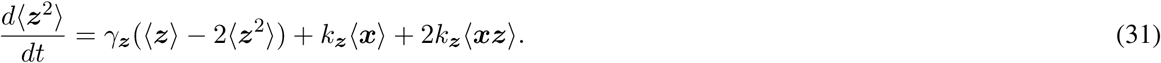

Steady-state analysis of this system of differential equations yields the following noise in the cell number

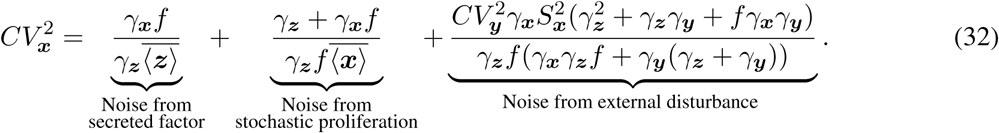

Comparing with (16) we see an additional third term that represents the contribution from the external disturbance, and this term monotonically decreases to zero as *f → ∞* (Fig. 3). Assuming a short-lived secreted factor *γ*_***z***_ ≫ *γ*_***x***_, *γ*_***y***_, (32) reduces to

**Figure 3:**
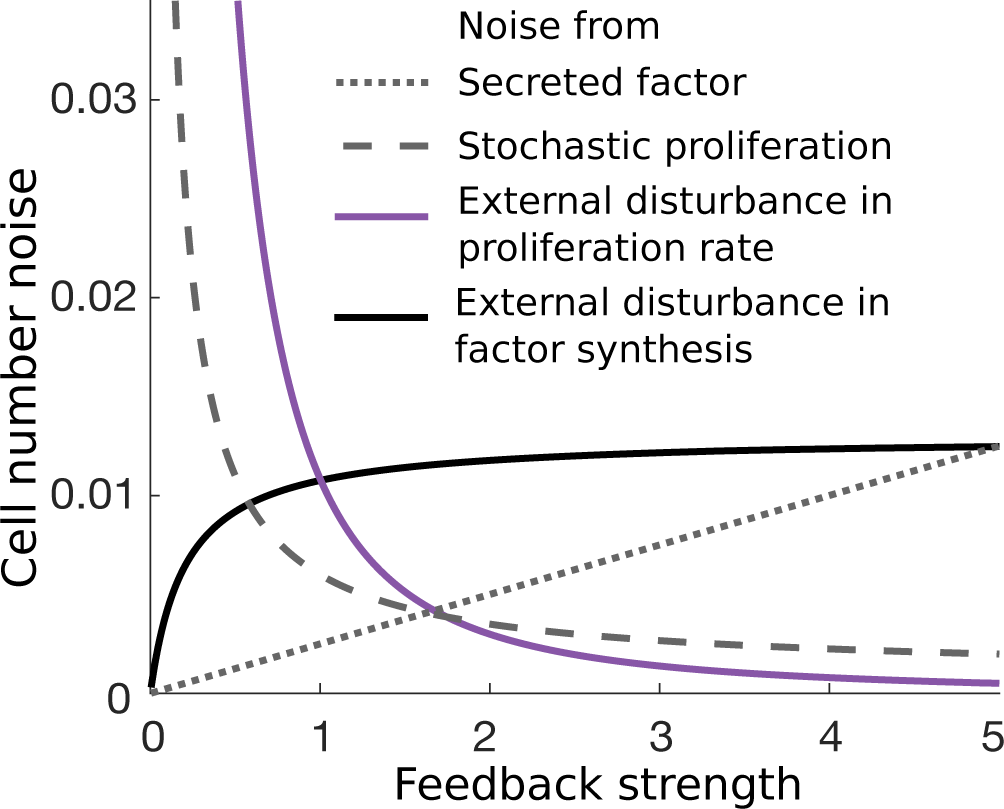
Depending on the source of noise, random fluctuations in cell numbers can be amplified or buffered with increasing negative feedback strength. Plots for the noise components shown in (32) and (35) as a function of *f*. While adding feedback buffers noise from stochastic proliferation and disturbances in the proliferation rate, it amplifies any noise associated with the secreted factor. The latter includes shot noise or Poisson fluctuation in secreted factor copy numbers, and external disturbances to the synthesis rate. Parameters chosen were: 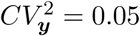, *γ*_*z*_ = 5*γ*_*x*_, *γ*_*y*_ = *γ*_*x*_/5, 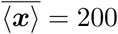, 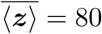.

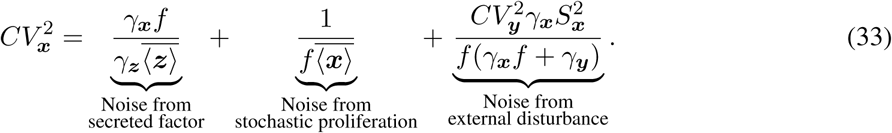

While the noise contribution from stochastic proliferation scales as *f*^−1^, depending on the relative values of *γ*_***x***_, *γ*_***y***_ the contribution from external disturbance scales between *f*^−1^ and *f*^−2^. This emphasizes a key point that feedback regulation is more efficient in suppressing noise arising from disturbances in proliferation rate as compared to the inherent stochasticity from small cell numbers. We next consider external disturbance in the synthesis rate *k*_***z***_ and revert the proliferation rate back to *g* (***z***).

### 2 External disturbance in the synthesis rate

The production and secretion of secreted factors is usually mediated by secondary molecules [58], and creates an alternative source of disturbance. To model this feature we modify the factor secretion rate as *k*_***z***_ (***y***), where ***y***(*t*) is the Ornstein-Uhlenbeck process (19), and as before, linearize the corresponding propensity function

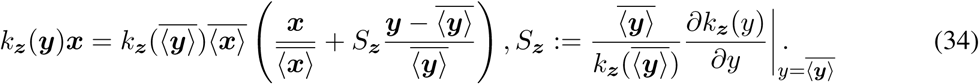

Here *S*_***z***_ is the log sensitivity of the secreted factor synthesis rate to the external disturbance. By performing an analysis similar to the previous section we obtain the following noise in cell numbers

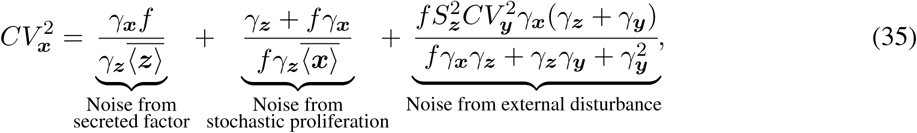

and now the contribution from external disturbance increases with increasing feedback strength *f* (Fig. 3). This is intuitive, as the disturbance acts through the secreted factor, and adding more feedback only functions to propagate this disturbance to affect cell numbers.

## 5 Conclusion

How cells regulate their population counts is an intriguing fundamental problem critical for functioning of cellular systems. For example in *Myxococcus xanthus* the formation of fruiting bodies only occurs in presence of precise cell densities and starvation. Further in *Dictyostelium*, during starvation, the cells aggregate to form spores in a fruiting body that is supported by a stalk [59]. If number of spores in the fruiting body are too small, then it will be too close to the ground and cause inefficient spore dispersal. However, if the number of spores is too large then it falls over, possibly ruining a chance of germination of the spores when nutrients become available. We systematically investigated the maintenance of precise cell numbers conferred by a negative feedback mechanism based on extracellular secretion of factors that are sensed by other cells in the population.

While producing low levels of secreted factors may be desirable to minimize energy costs and detection by predators, it comes at the cost of increased biomolecular noise in secreted factor copy numbers. Our analysis shows that while negative feedback suppresses cell number fluctuations from random birth/death of cells, it amplifies fluctuations arising from random birth/death of secreted factors (Fig 2). This results in cell number fluctuations being minimized at an intermediate feedback strength, and a fundamental limit to noise suppression given by (18). More specifically, the Fano factor (variance/mean or 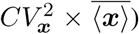 of the cell population count cannot be suppressed beyond 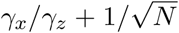. This implies that to have precise control, one not only needs a short-lived factor (*γ*_***x***_/*γ*_***z***_ ≪ 1) but also a high *N* (average number of factors secreted by an individual cell in its life span). Note that *γ*_***x***_/*γ*_***z***_ ≪ 1 and *N* = 4 gives a Fano factor of one, i.e., the number of cells will have a Poisson distribution, and much larger vales of *N* would be needed to obtain sub-Poisson statistics. Finally, we mention that similar results have been reported in the context of minimizing protein copy number fluctuations in auto-repressive genetic circuits, where noise sources are differentially effected by negative feedback creating optimal feedback strengths [60–65].

We also considered external disturbances in key model parameters, and depending on where the disturbance enters the systems, it can be amplified or buffered by negative feedback (Fig. 3). While in this work we have focused on a single cell type controlling its numbers, recent experiments reveal communication between different cell types via secreted growth factors to regulate population counts [66]. As part of our future work, we plan to extend the stochastic analysis to a system of multiple cell types interconnected by feedback loops.

## Acknowledgment

We thank Dr. Richard Gomer at the Texas A&M University for his helpful feedback on the manuscript. AS is supported by the National Institute of Health Grant 1R01GM126557-01.

